# Scoping review of the applications of peptide microarrays on the fight against human infections

**DOI:** 10.1101/2021.03.04.433859

**Authors:** Arthur Vengesai, Maritha Kasambala, Hamlet Mutandadzi, Tariro L. Mduluza-Jokonya, Takafira Mduluza, Thajasvarie Naicker

## Abstract

**Introduction:** This scoping review explores the use of peptide microarrays in the fight against infectious diseases. The research domains explored included the use of peptide microarrays in the mapping of linear B-cell and T cell epitopes, antimicrobial peptide discovery, immunosignature characterisation and disease immunodiagnostics. This review also provides a short overview of peptide microarray synthesis.

**Methods:** Electronic databases were systematically searched to identify relevant studies. The review was conducted using the Joanna Briggs Institute methodology for scoping reviews and data charting was performed using a predefined form. The results were reported by narrative synthesis in line with the Preferred Reporting Items for Systematic reviews and Meta-Analyses extension for Scoping Reviews guidelines.

**Results:** Eighty-six articles from 100 studies were included in the final data charting process. The majority (93%) of the articles were published during 2010–2020 and were mostly from Europe (44%) and North America (34 %). The findings were from the investigation of viral (44%), bacterial (30%), parasitic (25%) and fungal (2%) infections. Out of the serological studies, IgG was the most reported antibody type followed by IgM. The largest portion of the studies (78%) were related to mapping B-cell linear epitopes, 10% were on diagnostics, 9% reported on immunosignature characterisation and 6% reported on viral and bacterial cell binding assays. Two studies reported on T-cell epitope profiling.

**Conclusion:** The most important application of peptide microarrays was found to be B-cell epitope mapping or antibody profiling to identify diagnostic and vaccine targets. Immunosignatures identified by random peptide microarrays were found to be applied in the diagnosis of infections and interrogation of vaccine responses. The analysis of the interactions of random peptide microarrays with bacterial and viral cells using binding assays enabled the identification of antimicrobial peptides. Peptide microarray arrays were also used for T-cell linear epitope mapping which may provide more information for the design of peptide-based vaccines and for the development of diagnostic reagents.

## Introduction

Infectious diseases also known as communicable diseases are a major growing concern worldwide(1) and are a significant burden on public health (2). They account for a large proportion of death and disability globally. At least 25% of 60 million deaths that occur worldwide each year are estimated to be due to infectious diseases (2).

There are countless examples that highlight the severity of the impact of infectious diseases on human health (2). Since 31 December 2019 and as of 18 February 2021, **109 594 835** cases and almost 2.5 million deaths of COVID-19 have been reported world-wide (3). HIV infection continues to be a major pandemic where approximately 33 million people have died of HIV-related illnesses since the start of the pandemic. In 2019, 690 000 people died from HIV-related illnesses and 1.7 million people acquired new infections (4). Currently there are 20 Neglected tropical diseases’ (NTDs) affecting over 1.7 billion people and killing more than 200 000 people every year (5). Historically, the Black Death (1348–1350) killed 30%– 60 % of Europe’s population (2). In the 20th century, smallpox was responsible for an estimated 300–500 million deaths (2). The 1918-1919 Spanish Influenza pandemic killed more people than the World War 1 (2).

The threat posed by infectious diseases is further deepened by the continued emergence of new, unrecognized, and old infectious disease epidemics (2). Outbreaks caused by SARS-CoV-2, HIV, Ebola, influenza, and Zika viruses, have increased over the past decade, underlining the need for the rapid development of diagnostic tools and vaccines (6). **Fig. 1** illustrates epidemics that occurred in WHO African regions, during the period from 2016-2018, were 41countries (87%) had at least one epidemic, while 21 countries (45%) had at least one epidemic per year (7).

**Fig. 1:**
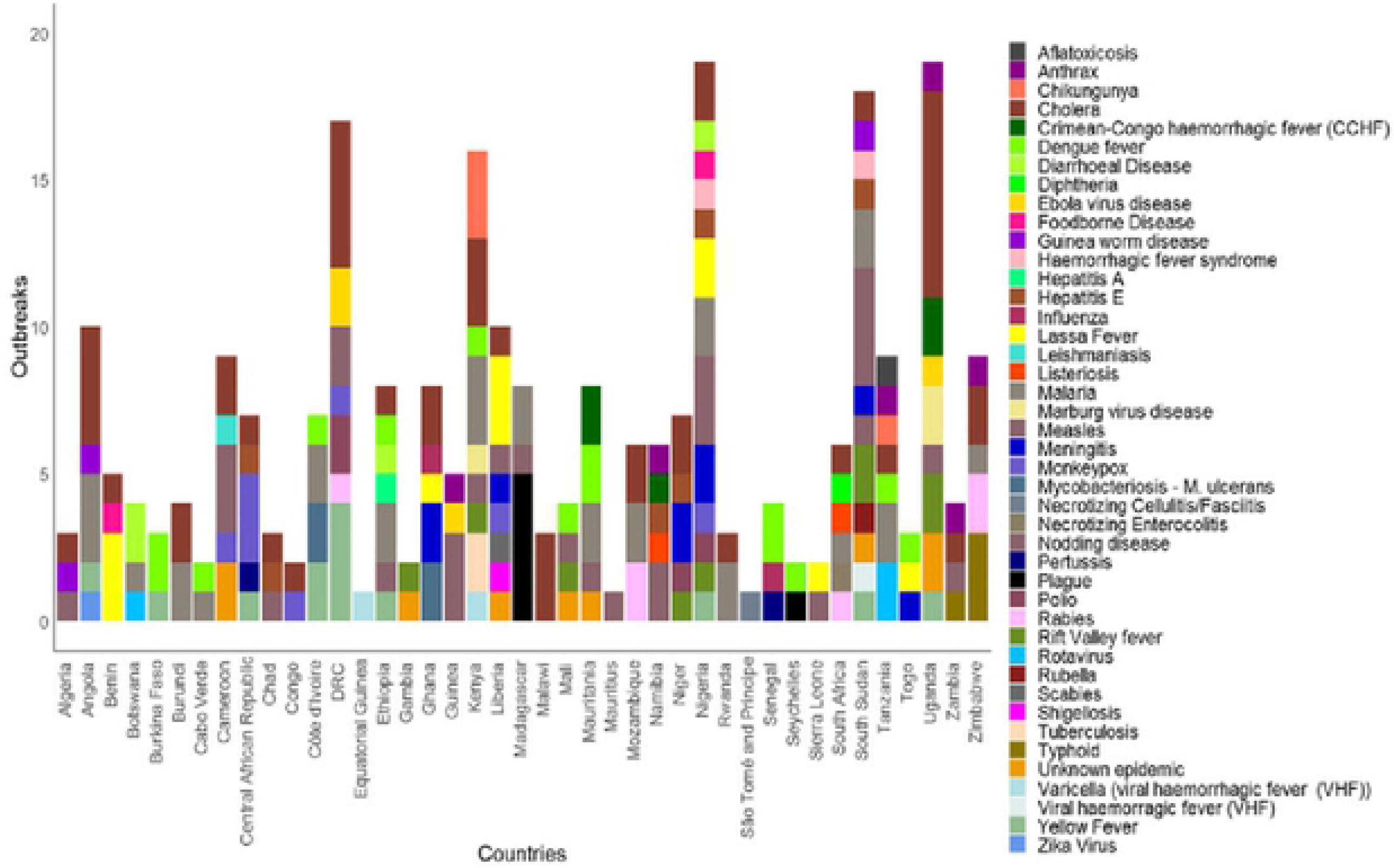
Infectious disease epidemics in the WHO African region, that occurred during the period 2016 to 2018.

A reasonable public health response towards addressing the infectious disease problem aims to address the fundamental factors that promote their occurrence and persistence, whilst implementing appropriate control measures (2). The field of medical biotechnology offers innovative devices for fighting infections, such as peptide microarrays (1,8).

Peptide microarrays are collections of short peptides immobilized on solid planar supports (9). They provide rapid, reproducible ways to simultaneously screen and detect hundreds to thousands of different pathogen related peptides or epitopes on standard microscope slides from small quantities of serum, plasma and cerebrospinal fluid (10–12). Peptide microarrays offer a wide range of applications in the fight against infectious diseases, such as, B-cell and T-cell epitope discovery for development of diagnostics and rationally designed vaccines, drug discovery (antimicrobial peptides discovery), immunosignature characterisation and pathogen immunodiagnostics (13–17). Additionally, peptide microarrays are used for autoimmune disease research, cancer research and enzyme profiling (18). In spite of the growing number of studies utilizing peptide microarrays, there is a paucity of systematic and narrative type reviews that reflect their clinical importance. This review focuses on the applications and use of peptide microarrays to fight infections

### Review aim and objectives

In order to systematically summarize the literature on the applications of peptide microarrays, we have conducted a scoping review. This scoping review aims to explore the use of peptide microarrays, in the mapping of B-cell linear epitopes, antimicrobial peptide discovery through bacterial cells glyco-profiling, immunosignature characterisation, immunodiagnostics and T-cell epitope mapping. This review also provides a short overview of peptide microarray synthesis. It is hoped that this review will highlight and enable recommendations that may aid future peptide microarray biomedical research, systematic and meta-analysis reviews.

## Methods

### Study design

The scoping review protocol was developed using the methodological framework proposed by Arksey and O’Malley (2005) and further refined by the Joanna Briggs Institute (19,20). The completed review followed the Preferred Reporting Items for Systematic reviews and Meta-Analyses extension for Scoping Reviews (PRISMA-ScR) guidelines (**S1 Table**) (21).

The review team consisted of four authors (AV, MK, HM and TLJ) who developed clear research questions, search strategies, identified relevant articles, selected articles, extracted and charted data. The discussion and reporting of the results were done in consultation with TN and TM.

### Eligibility criteria

The inclusion criteria was developed using the population-concept-context framework (19). The ‘population’ of the review were human participants of all ages, ethnicity and gender diagnosed with infectious diseases. Animal models for human infectious diseases studies, viruses and bacteria in antimicrobial activity investigations were also included as the review population. The ‘concept’ of the review was peptide microarrays. The review ‘context’ was a primary research study from any healthcare settings or institution from any country. All narrative reviews, studies investigating animal diseases, and duplicate articles were excluded. The search strategy was not restricted by the publication date or language. Hence, all related studies up to November 30, 2020, that met the inclusion criteria were assessed. As a scoping review is an iterative process, the eligibility criteria was amended as the study progressed.

### Search strategy information sources and search terms

The following online data bases PubMed, Medline complete, The Cochrane Central Register of Controlled Trials (CENTRAL) and Web of Science were systematically searched from their inception without any restrictions on language or date of publication. The data bases were searched using predefined keywords. Table 1 illustrates the search terms and strategy for PubMed which was adapted for the other databases. Additionally grey literature databases GreyLit and OpenGrey were searched and a manual search of the reference lists of relevant publications and reviews was conducted.

**Table 1:** Search Strategy in PubMed

| Search number | Query | Results |
| --- | --- | --- |
| 4 | ((("Peptides"[Mesh] OR Peptide*[tiab] OR Epitope*[tiab] AND (humans[Filter])) AND ("Microarray Analysis"[Mesh] OR Microarray*[tiab] OR Biochip*[tiab] OR Chip*[tiab] OR Array*[tiab] AND (humans[Filter]))) AND (Infestation*[tiab] OR Infection*[tiab] OR "Infectious disease*" [tiab] OR "Communicable disease*" [tiab] OR "Contagious disease*" [tiab] OR "Transmissible disease*" [tiab] OR Pathogen[tiab] OR Pathogens[tiab] AND (humans[Filter])) | 3,337 |
| 3 | Infestation*[tiab] OR Infection*[tiab] OR "Infectious disease*" [tiab] OR "Communicable disease*" [tiab] OR "Contagious disease*" [tiab] OR "Transmissible disease*" [tiab] OR Pathogen [tiab] OR Pathogens [tiab] | 1,098,661 |
| 2 | "Microarray Analysis" [Mesh] OR Microarray* [tiab] OR Biochip* [tiab] OR Chip* [tiab] OR Array* [tiab] | 178,487 |
| 1 | "Peptides" [Mesh] OR Peptide* [tiab] OR Epitope* [tiab] | 1,720,729 |

[Tiab] means the title and abstract were searched

### Review process and data charting

The retrieved literature were downloaded into Mendeley reference manager, and duplicates were removed. One reviewer (AV) assessed the titles of the studies identified by the search and excluded irrelevant studies. Two reviewers (AV and MK) independently assessed the eligibility of the abstracts and full texts of the retrieved studies to avoid bias. After the articles were selected, data was extracted and recorded in the excel spreadsheet. One author (AV) extracted and recorded the data from each study according to a pretested data extraction excel spreadsheet form (additional file 1) and a second reviewer (HM or MK or TMJ) verified the extracted data. Discrepancies were resolved by consensus and a third evaluator. The extracted data were author, date of publication, DOI, Aim and study domain, geographical location, microorganism or infection, antibody type, epitope prediction/selection, peptide synthesis, microarray printing and key findings.

### Methodological quality appraisal and analysis of the evidence

Methodological quality or risk of bias of the included articles was not appraised, which is consistent with guidance on scoping review conduct (19). The narrative synthesis of the results of this review were done in line with the recommendations set out in the PRISMA-ScR (21).

## Results and discussion

### Identification of potential studies

Electronic searches of seven databases yielded a total of 5929 articles (Pubmed: 3337, Medline (EBCOhost):1223, Cochrane: 17, Web of science: 1232, MedRxiv: 118, Greylit: 0, Open Grey: 2). Additional articles identified through manual searching yielded 11 articles that led to a total of 5940 titles and abstracts eligible for screening. A total of 253 full text articles were screened for eligibility after the removal of duplicate articles and irrelevant articles. Full text screening led to a total of 86 articles (100 studies) that were included in the scoping review. Two records were unable to be obtained in full-text format. **Fig. 2** is showing the flow chart of the studies identification and selection process.

**Fig 2:**
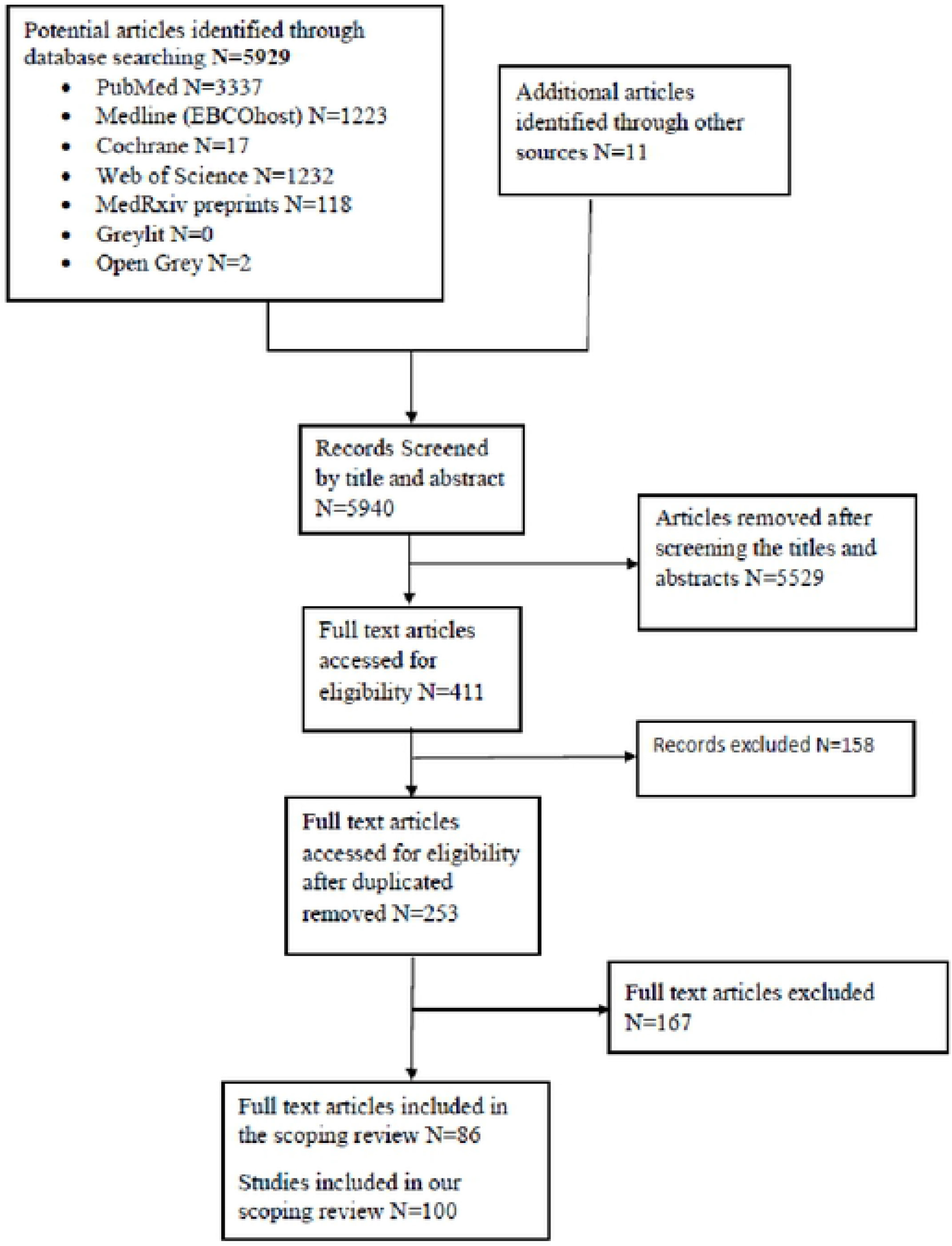
Flow chart of the studies identification and selection process

### Characteristics of the included articles

Characteristics of the included studies are shown in **S2 Table**. There were no articles published before 2001 on the study area and the peer-reviewed literature on the study area has increased considerably in the last few years (**Fig. 3**). Among the articles included, 63 % were published in the last five years (2015-2020) and approximately, 93% have been published in the last decade (2010–2020) of this current study.

**Fig 3:**
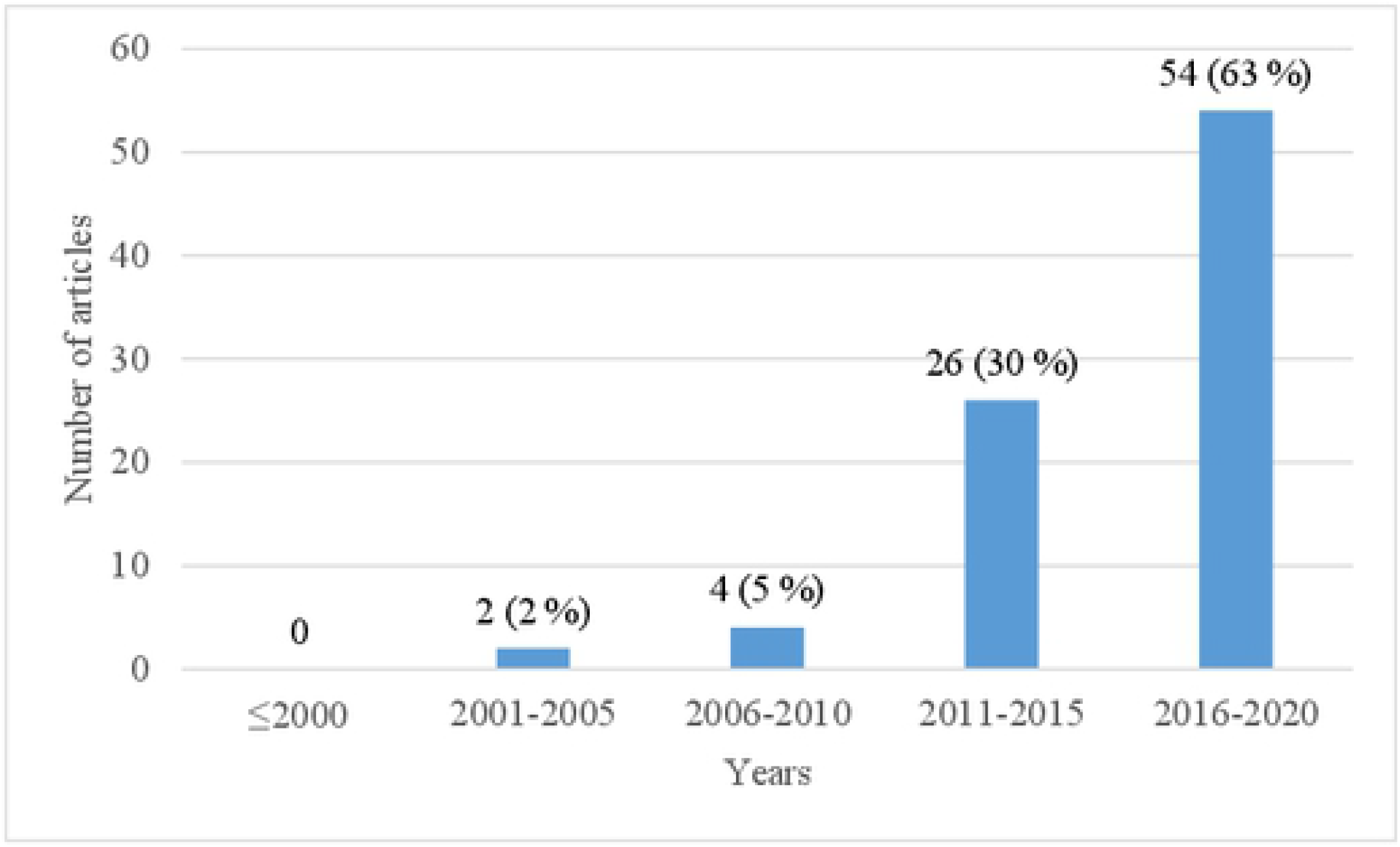
Number of included articles by year (2001-2020)

The included articles, were mainly from Europe 38 (44%) and North America 29 (34%) **Fig. 4**. From South America (Argentina 3 and Brazil 7) and Asia (China 7, Japan 1, and Sri Lanka 1) we included 10 (12%) and 9 (10%) articles respectively. Articles from Europe were divided among several countries, Germany 19, Sweden 6 Switzerland 2, Belgium 2, Denmark 2, Italy 4, Finland 1 Spain 1 and Austria 1. Articles from North America were mostly from USA 29, with one article from Cuba.

**Fig 4:**
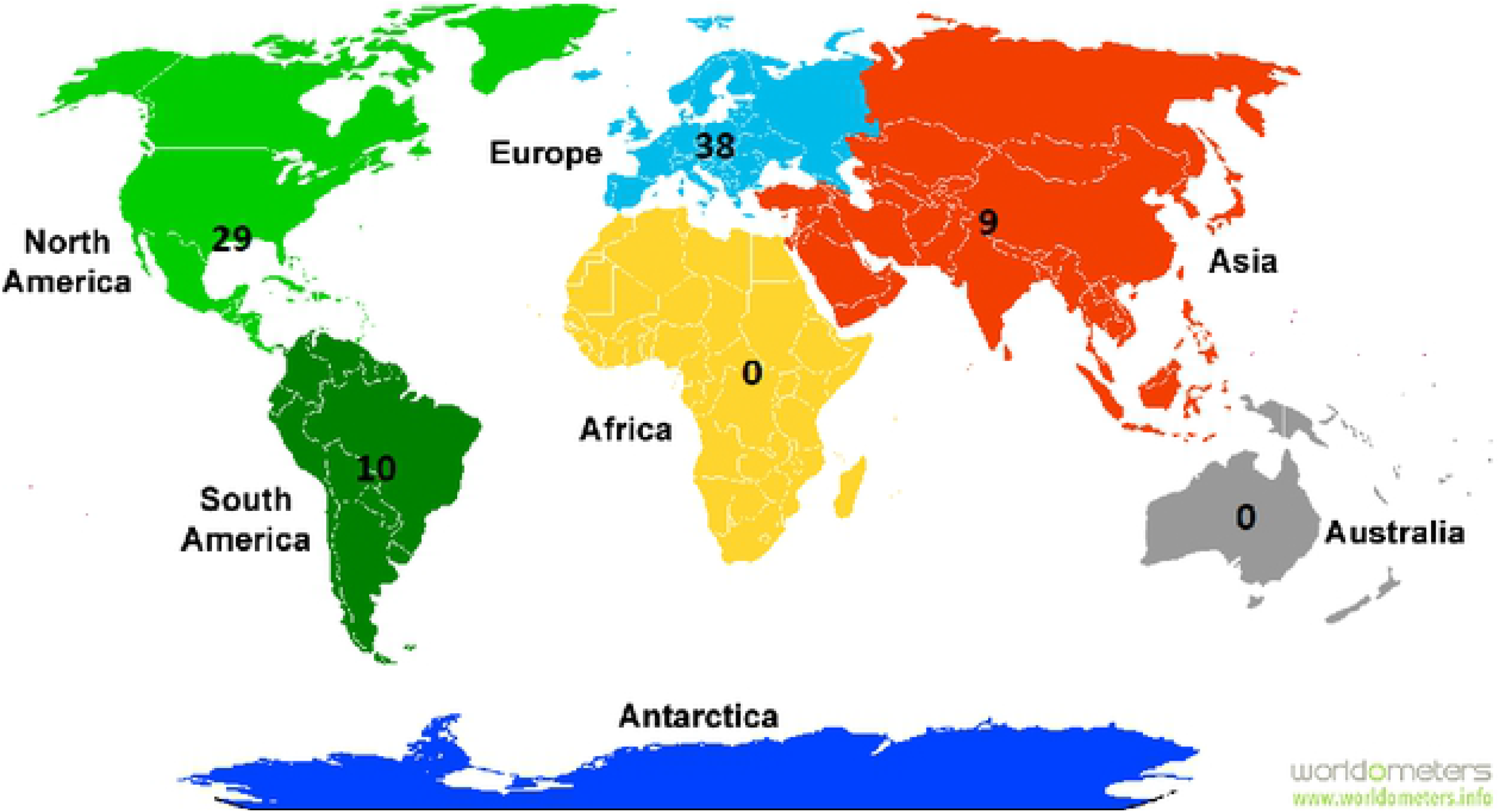
Number of articles included in the review by continent

In terms of the pathogens or infectious disease category studies (N=100), most studies were investigating viruses 43% (including SARS-CoV-2, HIV, Ebola) followed by studies investigating bacteria 30% (including TB, Lyme disease, chlamydia trachomatis) and Parasites 25% (Toxoplasma gondi, *S. mansoni, Plasmodium* species). Hitherto enigmatic diseases were investigated in 1 % of the included studies and Fungi (Coccidiodes) was investigated in 2 % of the studies. Two studies investigated health humans’ immunosignatures. Out of the 88 peptide microarray serological studies included IgG was the most invested antibody type followed by IgM. The IgG response shows a more specific binding pattern (less noise) than the IgM response, which reflects the higher specificity of IgGs (22). Two studies by Mishara *et al* (23) and Tokarz *et al* (24) investigated IgG and IgM profiles in cerebrospinal fluid.

### Peptide microarrays

#### PEPTIDE/EPITOPE IDENTIFICATION AND PREDICTION

B-cell and T-cell epitopes play a vital role in the development of peptide based vaccines and therapeutics and in the diagnosis of diseases (25,26). In this review, 6 methods were used for the identification and prediction of epitopes. These were computational overlapping peptides sequences, computational permutation scans, published synthetic peptides, computational random peptide sequences, phage display library and *in silico* prediction.

For epitope identification using overlapping peptides, the linear amino acid sequence of a protein is cut into peptides with overlapping sequences (27). This is achieved by shifting a frame of a distinct peptide length of a protein sequence of interest (28). In a permutation scan, each of the amino acid residues in a known antibody binding peptide is substituted by all amino acids or by one amino acid for example alanine permutation scans (29). Expect for *in silico* prediction methods, computational overlapping peptides sequences, computational permutation scans, computational random peptide sequences and phage display library peptide/epitope prediction methods are costly and time-consuming and demands large resources as they require screening of large arrays of potential epitope candidates. *In silico* prediction methods reduce the burden associated with epitope mapping by decreasing the list of potential epitope candidates for experimental testing(30,31). **Table 2** lists bioinformatics tools for the *in silico* prediction of epitopes on proteins for the studies included in this review. BepiPred 1.0 was the most frequently used software.

**Table 2:** B-cell epitope prediction software.

| Software | Server |
| --- | --- |
| MLCE | http://bioinf.uab.es/BEPPE |
| ABCpred | http://www.imtech.res.in/raghava/abcpred/ |
| BepiPred 1.0 | www.cbs.dtu.dk/services/BepiPred/ |
| Epitopia web server | http://epitopia.tau.ac.il/ |
| Antigenic | http://www.bioinformatics.nl/cgi-bin/emboss/antigenic |
| BCPREDS | http://ailab.ist.psu.edu/bcpred/ |
| Bcepred | http://crdd.osdd.net/raghava/bcepred/bcepred_instructions.html |

Peptide microarrays displayed short peptides (ranging 10–20 amino acid residues). Of note, most peptide microarrays displayed peptides with 15 amino acid residues, this length covers 83% of known linear antibody epitopes in the LANL immunology database, including the median length of epitopes (11 amino acids) (32). A few peptide microarrays displayed peptides with 5 and 6 amino acid residues set which are the shortest assumed B-cell epitope lengths (33).

#### PEPTIDE SYNTHESIS

Solid phase peptide synthesis (SPPS), was the method of choice for the production of peptides for most articles, although solution phase synthesis can still be useful for large-scale production of peptides. SPPS can be defined as a process in which a peptide anchored by its C-terminus to an insoluble polymer is assembled by the successive addition of protected amino acids constituting its sequence (34). SPPS, dramatically changed the strategy of peptide synthesis and simplified the tedious and demanding steps of purification associated with solution phase synthesis. SPPS also permitted the development of automation (35).

S synthesis is a special type of SPPS using cellulose as the solid support was used in 38 % of the studies that were included in the review. SPOT synthesis is a robust, rapid, and cost effective method for the simultaneous parallel chemical synthesis of peptides in a miniaturized array (36). SPOT synthesis has several advantages: cellulose is inexpensive and withstands the organic solvents and acids used during peptide synthesis. In addition, cellulose is stable in aqueous solutions and, because it is non-toxic, it is appropriate for screening biological samples. Another advantage of using SPOT synthesis on cellulose is the possibility of modifying the peptide (37). However, SPOT synthesis on porous membranes has its limitations when reducing the spot size below 1 mm and becomes costly and tedious when large numbers of copies of an identical array are required (38).

Peptide laser printing technology offered by PEPperPRINT Inc. (Heidelberg, Germany) (18) was used to produce peptides in 15 % of the included studies. The peptides are produced using a process based on electrostatic deposition and conjugation of dry amino acids, similar to the method used by laser printers.

#### PEPTIDE MICROARRAY SYNTHESIS

In general, two methods were used for the synthesis of peptide microarrays: the immobilization of pre-synthesized peptides and *in situ* synthesis of peptides on a solid support. Immobilization of pre-synthesised peptides involved SPOT synthesis, cleavage of solid phase bound peptides from the cellulose support matrix and spotting of the soluble peptides onto various types of planar surfaces for example glass chips using either a contact printer or a non-contact printer which minimizes contamination (39). Common solid phase materials such as functionalized polypropylene and glass were used for SPPS based *in-situ* peptide microarrays and cellulose was used for SPOT based *in situ* peptide microarray synthesis (40). The background signal from the *in situ* synthesis method is relatively lower than that produced by immobilizing pre-synthesized peptides because the background surface is selectively inert. However, the quality of peptides from the *in situ* synthesis method is lower than that of the spotting method because the peptide synthesized on a chip cannot be purified. Another problem yet to be solved with all *in situ* systems reported to date is the molecular characterization of the peptides. The lack of direct, *in situ* peptide analysis remains a major roadblock in the development of high-quality peptide arrays (40).

Peptide microarrays are offered by various providers, **Table 3** list the companies and the peptide microarray synthesis method including peptide synthesis and solid phase used by the companies. Peptide microarray providers are not limited to those included in **Table 3**. Of importance, Suzhou Epitope (Suzhou, China) uses polymer coated initiator integrated poly(dimethysiloxane) (iPDMS), as a solid supporting material. With an excellent capacity for preventing or reducing non-specific interactions, iPDMS, is able to provide near zero background for microarray screening. iPDMS can also achieve an extremely low limit of detection (41).

**Table 3:**
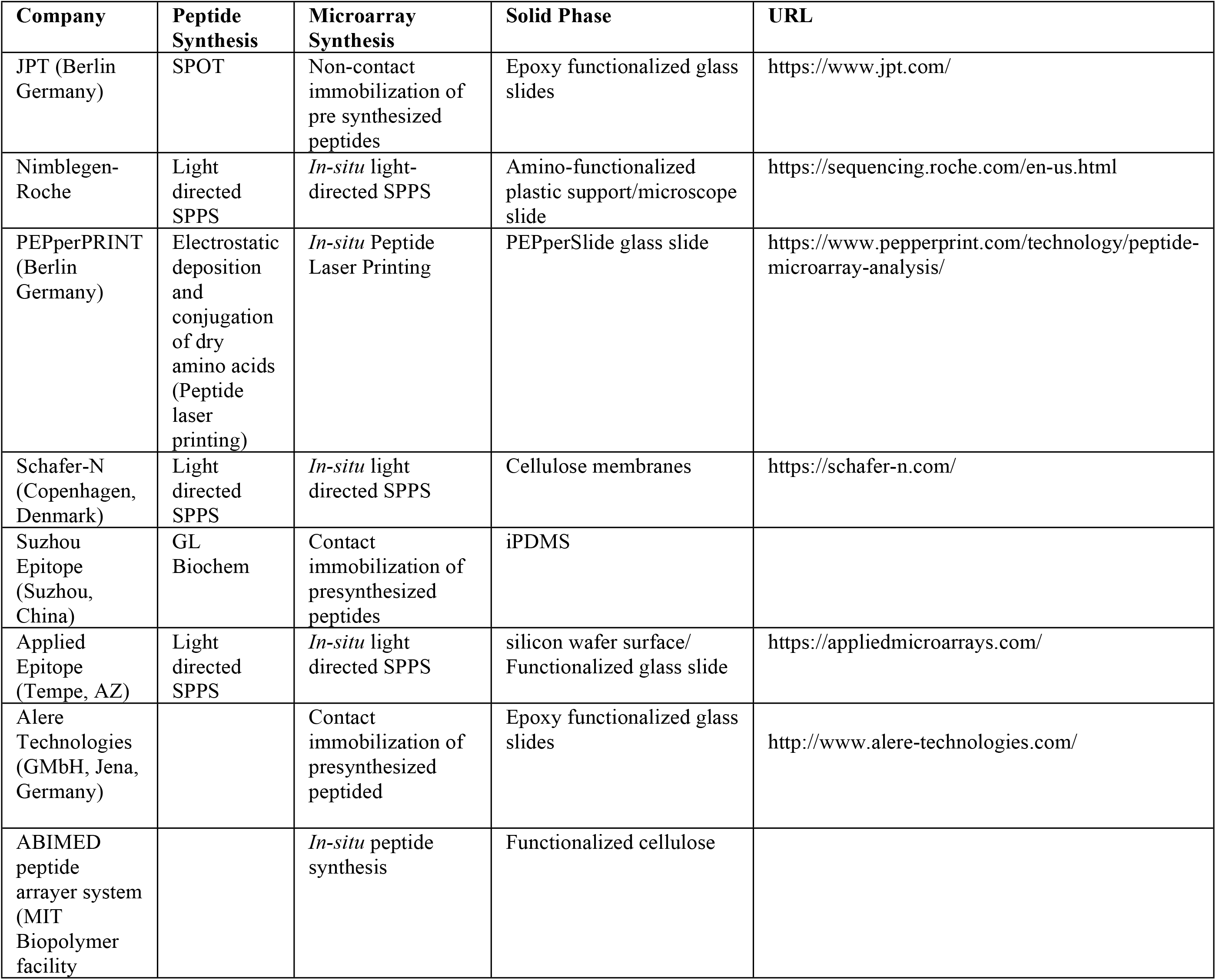
Peptide microarrays synthesis companies and the peptide microarray synthesis method including peptide synthesis and solid phase.

### Research domains

For the purpose of narrative review, based on the main research objectives, studies were classified into one of the following five research domains: mapping of B-cell linear epitopes, binding assays, immunosignatures characterisation, immunodiagnostics and mapping T-cell epitopes. The largest portion of the studies were related to mapping B-cell linear epitopes 78%, followed by studies on immunosignature characterisation 9%, while 8% reported immunodiagnostics and 6% reported virus and bacterial binding assays. Two studies reported mapping T-cell epitopes.

#### MAPPING B-CELL LINEAR EPITOPES

Antibodies recognize and bind their target protein antigens via surface accessible interaction sites, the linear epitopes or the conformational epitopes (38). High-content peptide microarrays allow linear epitope profiling of entire pathogen proteomes (42). There is great interest in identifying epitopes in antigens for a number of practical reasons(30,43). In the review, characterization of antibody specificities was mostly used to identify epitopes with potential applications in diagnosis of diseases. Epitope mapping identified epitopes useful in monitoring immune responses after chemotherapeutic treatments and vaccinations and for vaccine development. One study used epitope mapping to identify disease aetiology (44). Studies used overlapping peptides for the general epitope mapping and permutation scans or substitution analysis for fine epitope mapping. However it should be noted that mapping of B-cell epitopes using overlapping synthetic peptides permits the elucidation of linear epitopes only (45).

#### BACTERIA, VIRUS AND LIPOPOLYSACCHARIDES BINDING ASSAYS

Peptides can bind to various targets including bacterial and viral cells and lipopolysaccharides (LPS)(46). In the current review, peptide microarray binding assays were used to uncover the cyclic di-GMP (c-di-GMP) binding site of a *Pseudomonas aeruginosa* protein (PA3740), the Toll-like receptor (TLR) amino acid sequence for bacterial cell binding peptides and random peptide microarrays were used to screen for antimicrobial peptides (AMP).

The rise of multi-drug resistant pathogens is one of the most important global health issues and demands new compounds with novel mechanisms to combat these pathogens(47,48). However, new drug discovery has not kept pace as they often suffer with cross-resistance from existing agents of similar structure. Short, cationic peptides with antimicrobial activity known as AMPs, are essential to the host defences (48). AMPs are promising alternative to traditional antimicrobial drugs. AMPs are a diverse family of short peptides, between 5 and 50 amino acids in length and most possess an overall net positive charge to their structure (49) that display a broad spectrum killing properties to all pathogens. They are fast acting and have a decreased likelihood to induce pathogenic resistance as compared to traditional antimicrobial drugs and therefore could be next generation antibiotics (49). Screening for AMPs using peptide microarrays is a very convenient tool in the development of these drug candidates (28). In the current review, Svarovsky and Gonzalez-Moa, 2011 used fluorescently labelled bacteria and LPS to discover peptide sequences that not only specifically bound to LPS, but incidentally also inhibited bacterial cell growth (50). Betanzos *et al*., 2009 using luminescent LPS-quantum dots from O111:B4 and O55:B5 serotypes of *E. coli* revealed that peptides binding to *E. coli* LPS were highly enriched in aromatic and cationic amino acids and most inhibited growth (51). Johnston *et al*., 2017 screened a range of pathogens (10 viruses and 11 bacteria) against a library of 10,000 peptides to identify shared and specific pathogen binding peptides that were used for the development of a pathogen binding 100-peptide microarray (52).

TLRs are membrane bound-receptors responsible for recognizing pathogen associated molecular patterns and activation of the immune system. They specifically, recognize LPS, eliciting immune responses against invading bacteria (46). In the current review, a study by Tanaka *et al*., 2018 revealed several TLR4 peptides, including GRHIFWRR that demonstrated binding to *Escherichia coli* as well as LPS. These peptides exhibited a high proportion of arginine and lysine residues, positive charge, and low GRAVY value (hydrophilic) (46). Düvel *et al*., 2015, using fluorescence labelled c-di-GMP, showed that PA3740 octomer peptides bind c-di-GMP with high affinity and uncovered LKKALKKQTNLR to be a putative c-di-GMP binding motif. (53).

#### IMMUNOSIGNATURES

There is an increasing awareness that health care must move from post-symptomatic treatment to pre-symptomatic intervention (54). A universal system to diagnose disease, characterize infection or evaluate the response to a vaccine would be useful (55). An ideal system would allow regular monitoring of health status using circulating antibodies to report on health fluctuations. Random peptide microarrays can do this through antibody signatures (54). An immunosignature is a pattern of binding of serum antibodies to an array of thousands of random-sequence peptides in a broad and unbiased fashion (15,56).

Immunosignatures are not based on natural peptide sequence, but instead on a representative and diverse chemical space, a fact that simplifies peptide synthesis (57). Antibodies will bind to random peptides under permissive binding conditions. The binding is detected by a fluorescent anti-human secondary antibody. A high-resolution laser scanner provides an intensity value for each peptide (15). Querying immunosignature data using statistical and machine learning the random patterns of antibody peptide interactions can be used to diagnose disease, even many diseases simultaneously (15). In this review, this approach was shown to have diagnostic and prognostic potential for diseases and interrogation of vaccine response.

#### IMMUNODIAGNOSTICS

Serological assays play a major role in the diagnosis of both past and recent infections (24,58). These assays often based on crude antigen extracts or purified native antigenic proteins or recombinant antigens have constrains. Production of native antigens is limited, and the amounts are difficult to standardize. There is risk of contamination with proteins from organisms used in the production of recombinant antigens. Moreover, some recombinant antigens show lower reactivity than their corresponding native antigens, due to differences in protein folding that can result in altered epitope presentation. To avoid these limitations, several studies have shown that peptide microarrays can be used in serological assays to discriminate infected individuals from healthy individuals (16,24,58–60). Peptide microarray immunoassays were also shown to be capable of simultaneous multiplex diagnosis of different pathogens with a single patient serum sample (16,24,61). However, it is extremely unlikely that a single peptide can distinguish pathogens or strain types reliably (59). To achieve a satisfactory diagnostic sensitivity and a high specificity, it is necessary to use optimized peptide combinations, mimicking reactive epitopes on natural antigens. This strategy improves assay specificity by eliminating non-specific and potentially cross-reactive epitopes. Targeting a combination of such antigens can enhance assay sensitivity and has been shown to improve the diagnosis of tick-borne diseases (24). To select candidate diagnostic peptide sequences for subsequent analysis, *in-silico* predicted B-cell epitopes and previously predicted diagnostic peptides were used in the studies included in the review.

#### MAPPING T-CELL EPITOPES

Rational development and evaluation of peptide based vaccines and therapeutics requires identification and measurement of epitope-specific CD4 and CD8 T-cell responses. Conventional T-cell epitope discovery methods are labour intensive and do not scale well (62). In the current review, two studies (13,62) described the use of peptide microarrays using overlapping peptides, major histocompatibility complexes (MHC) and fluorescent tagged anti-MHC antibodies to map immunodominant T-cell epitopes. This high-throughput identification of T-cell epitopes will provide more information for the design of peptide-based vaccines and for the development of diagnostic reagents, such as MHC peptides.

### Strength and limitation

A clear limitation of conventional peptide microarrays is their restriction to linear protein epitopes, whereas conformational epitope antibody recognition cannot be identified (63). Detection of antibodies recognizing all potential epitopes whether linear, conformational or carbohydrate or LPS is a key requirement to comprehensively profile the humoral immune response (55).

The main advantage of the peptide microarray design is the miniaturisation of antibody-antigen interaction assays, the simultaneous analysis of several peptide sequences and the subsequent reduction in serum volume required from patients since this always represents a limiting factor in serological studies (64). By using peptide microarrays, it is feasible to simultaneously investigate the prevalence of the respective antibody classes in a specific patient and to differentiate the reactivity to all epitopes recognized by the different antibody class. By using different fluorescently labelled secondary antibodies each recognizing a particular antibody class, peptide microarrays permits the detection of different antibody classes within the same microarray (65).

In binding assays a distinct advantage offered by the peptide microarrays is the immediate visual assessment of all bacterial and viral cells and LPS binding events, enabling the parallel analysis of all binding peptides at once. This is useful for selection of orthogonal functional peptides that have different binding targets. A distinct disadvantage, however, is the limited number of potential binding ligands that generally does not allow meaningful selection of consensus sequences or binding motifs (66).

## Conclusion

In the review the peptide microarrays were shown to offer a wide range of applications, including, B-cell and T-cell epitope discovery for development of diagnostics and vaccines, serological diagnosis of viruses and bacteria as well as parasitic diseases pathogen and antimicrobial peptides discovery. Their most important was shown to be B-cell epitope mapping or antibody profiling to identify diagnostics and vaccine targets. Immunosignatures identified by random peptide microarrays were shown to be applied in the diagnosis of infections and interrogation of vaccine responses. Analysing the interactions of random peptide microarrays with bacterial and viral cells using binding assays enabled the identification of antimicrobial peptides. Peptide microarray arrays were also used for T-cell linear epitope mapping which may provide more information for the design of peptide-based vaccines and for the development of diagnostic reagents.

## Acknowledgements

The authors would like to acknowledge the valuable input of Professor Francisca Mutapi.

## Supporting information

S1 Table. PRISMA Extension for Scoping Reviews guidelines checklist.

S2 Table. General characteristics of the studies included in the scoping review.

## References

1. Afzal H, Zahid K, Ali Q, Sarwar K, Shakoor S, Nasir U, et al. Role of Biotechnology in Improving Human Health. J Mol Biomark Diagn. 2016;07(06).

2. Nii-Trebi NI. Emerging and Neglected Infectious Diseases: Insights, Advances, and Challenges [Internet]. Vol. 2017, BioMed Research International. Hindawi Limited; 2017 [cited 2021 Jan 14]. Available from: https://pubmed.ncbi.nlm.nih.gov/28286767/

3. Coronavirus disease (COVID-19) [Internet]. [cited 2021 Jan 14]. Available from: https://www.who.int/emergencies/diseases/novel-coronavirus-2019

4. HIV/AIDS [Internet]. [cited 2021 Jan 14]. Available from: https://www.who.int/news-room/fact-sheets/detail/hiv-aids

5. Neglected tropical diseases | Uniting to Combat NTDs [Internet]. [cited 2021 Jan 14]. Available from: https://unitingtocombatntds.org/ntds/

6. Heiss K, Heidepriem J, Fischer N, Weber LK, Dahlke C, Jaenisch T, et al. Rapid Response to Pandemic Threats: Immunogenic Epitope Detection of Pandemic Pathogens for Diagnostics and Vaccine Development Using Peptide Microarrays. Cite This J Proteome Res [Internet]. 2020;19:4339–54. Available from: https://dx.doi.org/10.1021/acs.jproteome.0c00484

7. Talisuna AO, Okiro EA, Yahaya AA, Stephen M, Bonkoungou B, Musa EO, et al. Spatial and temporal distribution of infectious disease epidemics, disasters and other potential public health emergencies in the World Health Organisation Africa region, 2016-2018. Global Health [Internet]. 2020 Jan 15 [cited 2021 Jan 14];16(1):9. Available from: https://globalizationandhealth.biomedcentral.com/articles/10.1186/s12992-019-0540-4

8. Pham P V. Medical biotechnology: Techniques and applications. In: Omics Technologies and Bio-engineering: Towards Improving Quality of Life. Elsevier Inc.; 2018. p. 449–69.

9. Zandian A, Forsström B, Häggmark-Månberg A, Schwenk JM, Uhlén M, Nilsson P, et al. Whole-Proteome Peptide Microarrays for Profiling Autoantibody Repertoires within Multiple Sclerosis and Narcolepsy. J Proteome Res [Internet]. 2017 Mar 3 [cited 2020 Feb 26];16(3):1300–14. Available from: http://www.ncbi.nlm.nih.gov/pubmed/28121444

10. Cretich M, Gori A, D’Annessa I, Chiari M, Colombo G. Peptides for Infectious Diseases: From Probe Design to Diagnostic Microarrays. Antibodies [Internet]. 2019 Mar 12 [cited 2021 Jan 14];8(1):23. Available from: https://www.mdpi.com/2073-4468/8/1/23

11. Yun SG, Jang JW, Lee JH, Lim CS, Kim J, Ki Y, et al. Evaluation of Novel Multiplex Antibody Kit for Human Immunodeficiency Virus 1/2 and Hepatitis C Virus Using Sol-Gel Based Microarray. Biomed Res Int [Internet]. 2015 [cited 2021 Jan 14];2015. Available from: https://pubmed.ncbi.nlm.nih.gov/26457305/

12. Mendes TA de O, Reis Cunha JL, de Almeida Lourdes R, Rodrigues Luiz GF, Lemos LD, dos Santos ARR, et al. Identification of Strain-Specific B-cell Epitopes in Trypanosoma cruzi Using Genome-Scale Epitope Prediction and High-Throughput Immunoscreening with Peptide Arrays. Marques ETA, editor. PLoS Negl Trop Dis [Internet]. 2013 Oct 31 [cited 2021 Jan 14];7(10):e2524. Available from: https://dx.plos.org/10.1371/journal.pntd.0002524

13. Malnati MS, Heltai S, Cosma A, Reitmeir P, Allgayer S, Glashoff RH, et al. A new antigen scanning strategy for monitoring HIV-1 specific T-cell immune responses. J Immunol Methods [Internet]. 2012 Jan 31 [cited 2020 Nov 23];375(1–2):46–56. Available from: https://pubmed-ncbi-nlm-nih-gov.ukzn.idm.oclc.org/21963950/

14. Sabalza M, Barber CA, Abrams WR, Montagna R, Malamud D. Zika virus specific diagnostic epitope discovery. J Vis Exp [Internet]. 2017 Dec 12 [cited 2020 Nov 24];2017(130). Available from: https://pubmed-ncbi-nlm-nih-gov.ukzn.idm.oclc.org/29286404/

15. Stafford P, Wrapp D, Johnston SA. General assessment of humoral activity in healthy humans. Mol Cell Proteomics [Internet]. 2016 May 1 [cited 2020 Nov 24];15(5):1610–21. Available from: https://pubmed-ncbi-nlm-nih-gov.ukzn.idm.oclc.org/26902205/

16. Maksimov P, Zerweck J, Maksimov A, Hotop A, Groß U, Pleyer U, et al. Peptide microarray analysis of in silico-predicted epitopes for serological diagnosis of Toxoplasma gondii infection in humans. Clin Vaccine Immunol [Internet]. 2012 Jun [cited 2020 Nov 22];19(6):865–74. Available from: https://pubmed-ncbi-nlm-nih-gov.ukzn.idm.oclc.org/22496494/

17. Lee JS, Song JJ, Deaton R, Kim JW. Assessing the detection capacity of microarrays as bio/nanosensing platforms. Biomed Res Int [Internet]. 2013 [cited 2020 Nov 22];2013. Available from: https://pubmed-ncbi-nlm-nih-gov.ukzn.idm.oclc.org/24324959/

18. PEPperPRINT: Peptide Microarray Analysis [Internet]. [cited 2021 Jan 14]. Available from: https://www.pepperprint.com/technology/peptide-microarray-analysis/

19. Peters MDJ, Godfrey CM, Khalil H, McInerney P, Parker D, Soares CB. Guidance for conducting systematic scoping reviews. Int J Evid Based Healthc [Internet]. 2015 Sep 1 [cited 2021 Jan 14];13(3):141–6. Available from: https://pubmed.ncbi.nlm.nih.gov/26134548/

20. Arksey H, O’Malley L. Scoping studies: Towards a methodological framework. Int J Soc Res Methodol Theory Pract [Internet]. 2005 Feb [cited 2021 Jan 14];8(1):19–32. Available from: https://www.tandfonline.com/doi/abs/10.1080/1364557032000119616

21. Tricco AC, Lillie E, Zarin W, O’Brien KK, Colquhoun H, Levac D, et al. PRISMA extension for scoping reviews (PRISMA-ScR): Checklist and explanation [Internet]. Vol. 169, Annals of Internal Medicine. American College of Physicians; 2018 [cited 2021 Jan 14]. p. 467–73. Available from: https://pubmed.ncbi.nlm.nih.gov/30178033/

22. Heidepriem J, Krähling V, Dahlke C, Wolf T, Klein F, Addo MM, et al. Epitopes of Naturally Acquired and Vaccine-Induced Anti-Ebola Virus Glycoprotein Antibodies in Single Amino Acid Resolution. Biotechnol J. 2020 Sep 1;15(9).

23. Mishra N, Caciula A, Price A, Thakkar R, Ng J, Chauhan L V., et al. Diagnosis of Zika virus infection by peptide array and enzyme-linked immunosorbent assay. MBio. 2018 Mar 1;9(2).

24. Tokarz R, Mishra N, Tagliafierro T, Sameroff S, Caciula A, Chauhan L, et al. A multiplex serologic platform for diagnosis of tick-borne diseases. Sci Rep [Internet]. 2018 Dec 1 [cited 2020 Nov 23];8(1). Available from: https://pubmed-ncbi-nlm-nih-gov.ukzn.idm.oclc.org/29453420/

25. Saha S, Raghava GPS. Prediction of continuous B-cell epitopes in an antigen using recurrent neural network. Proteins Struct Funct Bioinforma [Internet]. 2006 Aug 7 [cited 2021 Feb 9];65(1):40–8. Available from: http://doi.wiley.com/10.1002/prot.21078

26. Li Pira G, Ivaldi F, Bottone L, Manca F. High throughput functional microdissection of pathogen-specific T-cell immunity using antigen and lymphocyte arrays. J Immunol Methods [Internet]. 2007 Sep 30 [cited 2021 Feb 9];326(1–2):22–32. Available from: https://pubmed.ncbi.nlm.nih.gov/17673252/

27. Weber LK, Palermo A, Kügler J, Armant O, Isse A, Rentschler S, et al. Single amino acid fingerprinting of the human antibody repertoire with high density peptide arrays. J Immunol Methods [Internet]. 2017 Apr 1 [cited 2020 Nov 24];443:45–54. Available from: https://pubmed-ncbi-nlm-nih-gov.ukzn.idm.oclc.org/28167275/

28. Winkler DFH, Campbell WD. The spot technique the spot technique: Synthesis and screening of peptide macroarrays on cellulose membranes. Methods Mol Biol. 2008;494:47–70.

29. Lagatie O, Van Dorst B, Stuyver LJ. Identification of three immunodominant motifs with atypical isotype profile scattered over the Onchocerca volvulus proteome. PLoS Negl Trop Dis [Internet]. 2017 Jan 26 [cited 2020 Nov 23];11(1). Available from: https://pubmed-ncbi-nlm-nih-gov.ukzn.idm.oclc.org/28125577/

30. Sanchez-Trincado JL, Gomez-Perosanz M, Reche PA. Fundamentals and Methods for T- and B-Cell Epitope Prediction. Vol. 2017, Journal of Immunology Research. Hindawi Limited; 2017.

31. Sanchez-Lockhart M, Reyes DS, Gonzalez JC, Garcia KY, Villa EC, Pfeffer BP, et al. Qualitative Profiling of the Humoral Immune Response Elicited by rVSV-ΔG-EBOV-GP Using a Systems Serology Assay, Domain Programmable Arrays. Cell Rep [Internet]. 2018 Jul 24 [cited 2021 Feb 9];24(4):1050–1059.e5. Available from: https://doi.org/10.1016/j.celrep.2018.06.077

32. Stephenson KE, Neubauer GH, Reimer U, Pawlowski N, Knaute T, Zerweck J, et al. Quantification of the epitope diversity of HIV-1-specific binding antibodies by peptide microarrays for global HIV-1 vaccine development. J Immunol Methods. 2015 Jan 1;416:105–23.

33. Jaenisch T, Heiss K, Fischer N, Geiger C, Bischoff FR, Moldenhauer G, et al. High-density peptide arrays help to identify linear immunogenic B-cell epitopes in individuals naturally exposed to malaria infection. Mol Cell Proteomics [Internet]. 2019 [cited 2020 Nov 24];18(4):642–56. Available from: https://pubmed-ncbi-nlm-nih-gov.ukzn.idm.oclc.org/30630936/

34. Al-Warhi TI, Al-Hazimi HMA, El-Faham A. Recent development in peptide coupling reagents. Vol. 16, Journal of Saudi Chemical Society. Elsevier; 2012. p. 97–116.

35. Amblard M, Fehrentz JA, Martinez J, Subra G. Methods and protocols of modern solid phase peptide synthesis [Internet]. Vol. 33, Molecular Biotechnology. Springer; 2006 [cited 2021 Jan 14]. p. 239–54. Available from: https://link.springer.com/article/10.1385/MB:33:3:239

36. Fraczyk J, Walczak M, Kaminski ZJ. New methodology for automated SPOT synthesis of peptides on cellulose using 1,3,5-triazine derivatives as linkers and as coupling reagents. J Pept Sci [Internet]. 2018 Feb 1 [cited 2021 Jan 14];24(2):e3063. Available from: http://doi.wiley.com/10.1002/psc.3063

37. Winkler DFH, Hilpert K, Brandt O, Hancock REW. Synthesis of peptide arrays using SPOT-technology and the CelluSpots-method. Methods Mol Biol [Internet]. 2009 [cited 2021 Jan 14];570:157–74. Available from: https://pubmed.ncbi.nlm.nih.gov/19649591/

38. Beutling U, Frank R. Epitope Analysis Using Synthetic Peptide Repertoires Prepared by SPOT Synthesis Technology. In: Antibody Engineering [Internet]. Berlin, Heidelberg: Springer Berlin Heidelberg; 2010 [cited 2021 Jan 14]. p. 537–71. Available from: http://link.springer.com/10.1007/978-3-642-01144-3_35

39. Dikmans A, Beutling U, Schmeisser E, Thiele S, Frank R. SC2: A novel process for manufacturing multipurpose high-density chemical microarrays. QSAR Comb Sci [Internet]. 2006 Nov [cited 2021 Jan 14];25(11):1069–80. Available from: http://doi.wiley.com/10.1002/qsar.200640130

40. F.H. Winkler D. Chemistry of SPOT Synthesis for the Preparation of Peptide Macroarrays on Cellulose Membranes. Mini Rev Org Chem [Internet]. 2011 Mar 28 [cited 2021 Jan 14];8(2):114–20. Available from: http://www.eurekaselect.com/openurl/content.php?genre=article&issn=1570-193X&volume=8&issue=2&spage=114

41. Huang M, Ma Q, Liu X, Li B, Ma H. Initiator Integrated Poly(dimethysiloxane)-Based Microarray as a Tool for Revealing the Relationship between Nonspecific Interactions and Irreproducibility. Anal Chem [Internet]. 2015 Jul 21 [cited 2021 Jan 14];87(14):7085–91. Available from: https://pubs.acs.org/doi/abs/10.1021/acs.analchem.5b00694

42. Pérez-Bercoff L, Valentini D, Gaseitsiwe S, Mahdavifar S, Schutkowski M, Poiret T, et al. Whole CMV proteome pattern recognition analysis after HSCT identifies unique epitope targets associated with the CMV status. PLoS One [Internet]. 2014 Apr 16 [cited 2020 Nov 22];9(4). Available from: https://pubmed-ncbi-nlm-nih-gov.ukzn.idm.oclc.org/24740411/

43. Carmona SJ, Nielsen M, Schafer-Nielsen C, Mucci J, Altcheh J, Balouz V, et al. Towards high-throughput immunomics for infectious diseases: Use of next-generation peptide microarrays for rapid discovery and mapping of antigenic determinants. Mol Cell Proteomics [Internet]. 2015 Jul 1 [cited 2020 Nov 22];14(7):1871–84. Available from: https://pubmed-ncbi-nlm-nih-gov.ukzn.idm.oclc.org/25922409/

44. Ferrara G, Valentini D, Rao M, Wahlström J, Grunewald J, Larsson LO, et al. Humoral immune profiling of mycobacterial antigen recognition in sarcoidosis and Löfgren’s syndrome using high-content peptide microarrays. Vol. 56, International Journal of Infectious Diseases. Elsevier B.V.; 2017. p. 167–75.

45. Torréns I, Reyes O, Ojalvo AG, Seralena A, Chinea G, Cruz LJ, et al. Mapping of the antigenic regions of streptokinase in humans after streptokinase therapy. Biochem Biophys Res Commun [Internet]. 1999 May 27 [cited 2021 Feb 9];259(1):162–8. Available from: https://pubmed.ncbi.nlm.nih.gov/10334933/

46. Tanaka M, Harlisa IH, Takahashi Y, Ikhsan NA, Okochi M. Screening of bacteria-binding peptides and one-pot ZnO surface modification for bacterial cell entrapment. RSC Adv. 2018;8(16):8795–9.

47. Bluhm MEC, Knappe D, Hoffmann R. Structure-activity relationship study using peptide arrays to optimize Api137 for an increased antimicrobial activity against Pseudomonas aeruginosa. Eur J Med Chem [Internet]. 2015 Oct 20 [cited 2020 Nov 22];103:574–82. Available from: https://pubmed-ncbi-nlm-nih-gov.ukzn.idm.oclc.org/26408816/

48. Fjell CD, Jenssen H, Hilpert K, Cheung WA, Panté N, Hancock REW, et al. Identification of novel antibacterial peptides by chemoinformatics and machine learning. J Med Chem. 2009 Apr 9;52(7):2006–15.

49. López-Pérez PM, Grimsey E, Bourne L, Mikut R, Hilpert K. Screening and optimizing antimicrobial peptides by using SPOT-synthesis [Internet]. Vol. 5, Frontiers in Chemistry. Frontiers Media S. A; 2017 [cited 2021 Jan 14]. p. 25. Available from: www.frontiersin.org

50. Svarovsky SA, Gonzalez-Moa MJ. High-throughput platform for rapid deployment of antimicrobial agents. ACS Comb Sci. 2011 Nov 14;13(6):634–8.

51. Betanzos CM, Gonzalez-Moa MJ, Boltz KW, Vander Werf BD, Johnston SA, Svarovsky SA. Bacterial glycoprofiling by using random sequence peptide microarrays. ChemBioChem [Internet]. 2009 Mar 23 [cited 2021 Jan 14];10(5):877–88. Available from: http://doi.wiley.com/10.1002/cbic.200800716

52. Johnston SA, Domenyuk V, Gupta N, Batista MT, Lainson JC, Zhao ZG, et al. A Simple Platform for the Rapid Development of Antimicrobials. Sci Rep. 2017 Dec 1;7(1).

53. Düvel J, Bense S, Möller S, Bertinetti D, Schwede F, Morr M, et al. Application of synthetic peptide arrays to uncover cyclic di-GMP binding motifs. J Bacteriol. 2016;198(1):138–46.

54. Legutki JB, Zhao ZG, Greving M, Woodbury N, Johnston SA, Stafford P. Scalable high-density peptide arrays for comprehensive health monitoring. Nat Commun. 2014 Sep 3;5.

55. Legutki JB, Magee DM, Stafford P, Johnston SA. A general method for characterization of humoral immunity induced by a vaccine or infection. Vaccine [Internet]. 2010 Jun 17 [cited 2020 Nov 23];28(28):4529–37. Available from: https://pubmed-ncbi-nlm-nih-gov.ukzn.idm.oclc.org/20450869/

56. Navalkar KA, Johnston SA, Woodbury N, Galgiani JN, Magee DM, Chicacz Z, et al. Application of immunosignatures for diagnosis of valley fever. Clin Vaccine Immunol [Internet]. 2014 [cited 2020 Nov 23];21(8):1169–77. Available from: https://pubmed-ncbi-nlm-nih-gov.ukzn.idm.oclc.org/24964807/

57. Singh S, Stafford P, Schlauch KA, Tillett RR, Gollery M, Johnston SA, et al. Humoral immunity profiling of subjects with myalgic encephalomyelitis using a random peptide microarray differentiates cases from controls with high specificity and sensitivity. Mol Neurobiol [Internet]. 2018 Jan 1 [cited 2020 Nov 24];55(1):633–41. Available from: https://pubmed-ncbi-nlm-nih-gov.ukzn.idm.oclc.org/27981498/

58. Rizwan M, Rönnberg B, Cistjakovs M, Lundkvist Å, Pipkorn R, Blomberg J. Serology in the Digital Age: Using Long Synthetic Peptides Created from Nucleic Acid Sequences as Antigens in Microarrays. Microarrays. 2016 Aug 10;5(3):22.

59. Arranz-Solís D, Cordeiro C, Young LH, Dardé ML, Commodaro AG, Grigg ME, et al. Serotyping of Toxoplasma gondii Infection Using Peptide Membrane Arrays. Front Cell Infect Microbiol [Internet]. 2019 Nov 29 [cited 2020 Nov 24];9. Available from: https://pubmed-ncbi-nlm-nih-gov.ukzn.idm.oclc.org/31850240/

60. Bergamaschi G, Fassi EMA, Romanato A, D’Annessa I, Odinolfi MT, Brambilla D, et al. Computational analysis of dengue virus envelope protein (E) reveals an epitope with flavivirus immunodiagnostic potential in peptide microarrays. Int J Mol Sci [Internet]. 2019 Apr 2 [cited 2020 Nov 23];20(8). Available from: https://pubmed-ncbi-nlm-nih-gov.ukzn.idm.oclc.org/31003530/

61. Sachse K, Rahman KS, Schnee C, Müller E, Peisker M, Schumacher T, et al. A novel synthetic peptide microarray assay detects Chlamydia species-specific antibodies in animal and human sera. Sci Rep [Internet]. 2018 Dec 1 [cited 2020 Nov 24];8(1). Available from: https://pubmed-ncbi-nlm-nih-gov.ukzn.idm.oclc.org/29549361/

62. Haj AK, Breitbach ME, Baker DA, Mohns MS, Moreno GK, Wilson NA, et al. High-Throughput Identification of MHC Class I Binding Peptides Using an Ultradense Peptide Array. J Immunol. 2020 Mar 15;204(6):1689–96.

63. Loeffler FF, Pfeil J, Heiss K. High-density peptide arrays for malaria vaccine development. In: Methods in Molecular Biology [Internet]. Humana Press Inc.; 2016 [cited 2021 Jan 14]. p. 569–82. Available from: https://link.springer.com/protocol/10.1007/978-1-4939-3387-7_32

64. Fernández L, Bleda MJ, Gómara MJ, Haro I. Design and application of GB virus C (GBV-C) peptide microarrays for diagnosis of GBV-C/HIV-1 co-infection. Anal Bioanal Chem [Internet]. 2013 [cited 2020 Nov 23];405(12):3973–82. Available from: https://pubmed-ncbi-nlm-nih-gov.ukzn.idm.oclc.org/23232955/

65. The distribution and functions of immunoglobulin isotypes - Immunobiology - NCBI Bookshelf [Internet]. [cited 2021 Jan 14]. Available from: https://www.ncbi.nlm.nih.gov/books/NBK27162/

66. Svarovsky SA, Gonzalez-Moa MJ. High-throughput platform for rapid deployment of antimicrobial agents. ACS Comb Sci [Internet]. 2011 Nov 14 [cited 2021 Jan 14];13(6):634–8. Available from: https://pubs.acs.org/doi/abs/10.1021/co200088c

